# MSABrowser: dynamic and fast visualization of sequence alignments, variations, and annotations

**DOI:** 10.1101/2021.04.05.426321

**Authors:** Furkan M. Torun, Halil I. Bilgin, Oktay I. Kaplan

**Author notes:** Both authors contributed equally to this work (Furkan M. Torun, Halil I. Bilgin).

## Abstract

Sequence alignment is an excellent way to visualize the similarities and differences between DNA, RNA, or protein sequences, yet it is currently difficult to jointly view sequence alignment data with genetic variations, modifications such as post-translational modifications, and annotations (i.e. protein domains). Here, we develop the MSABrowser tool that makes it easy to co-visualize genetic variations, modifications, and annotations on the respective positions of amino acids or nucleotides in pairwise or multiple sequence alignments. MSABrowser is developed entirely in JavaScript and works on any modern web browser at any platform, including Linux, Mac OS X, and Windows systems without any installation. MSABrowser is also freely available for the benefit of the scientific community.

**Availability and implementation:** MSABrowser is released as open-source and web-based software under GNU General Public License, version 3.0 (GPLv3). The visualizer, documentation, all source codes, and examples are available at http://thekaplanlab.github.io/ and GitHub repository https://github.com/thekaplanlab/msabrowser.

**Supplementary information:** Supplementary data are available online.

## 1 Introduction

The next-generation sequencing (NGS) technologies have revolutionized the genomics field, thus revealing more than 700 million genetic variations in the human genomes and millions of genetic variants in non-human primates (Taliun *et al*., 2019; Karczewski *et al*., 2020; Sundaram *et al*., 2018; Locke *et al*., 2011; Rhesus Macaque Genome Sequencing and Analysis Consortium *et al*., 2007; The Marmoset Genome Sequencing and Analysis Consortium, 2014; Sherry *et al*., 1999). Furthermore, clinical scientists and researchers have identified thousands of variants associated with health and diseases. Additionally, genome-wide association studies (GWAS) systematically identified candidate genomic regions responsible for phenotypic differences (Ozaki *et al*., 2002; Landrum *et al*., 2020). All these data suggest that each genomic or proteomic position has a variety of unique details, including mutation, single-nucleotide polymorphism (SNP), orthologous variants, allele frequency, disease associations, DNA methylation, and amino acid phosphorylation at specific positions. Although substantial progress has been made incomparably visualizing specific positions on pair sequence alignment (PSA) and multiple sequence alignment (MSA) between humans and non-human species, annotating specific information to each position at DNA, RNA or protein sequences involves manual inspection with the incorporation of position-specific data. Challenges remain to co-visualize ever-increasing variants comparably along with variant-specific annotations. It is hence all-important that the variant visualization tool enables flexibility of annotating specific functional data to variants.

Here we, therefore, develop a free, open-source, user-friendly web-based tool called MSABrowser to dynamically and rapidly visualize multiple sequence alignments, with the integration of variant-specific annotations to the corresponding positions (**Fig. 1**). MSABrowser is based on a JavaScript programming language that enables users to construct interactive pages with complex features, so it works easily without installation on any modern web browser.

**Fig. 1.**
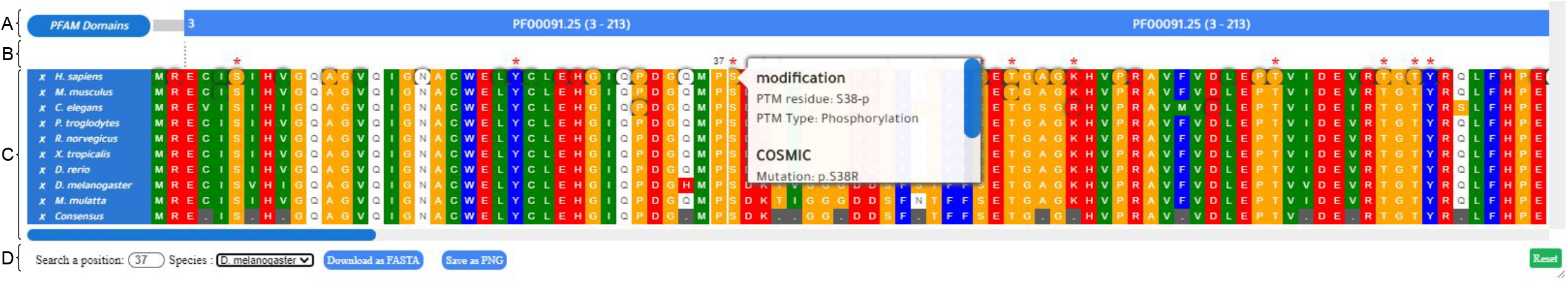
An overview of the MSABrowser tool. In this figure, MSA for homologous proteins of human TUBA1A protein is visualized together with genetic variations on the corresponding positions on sequences, and associated intervals such as protein domains are specified. **(A)** The annotation part represents the specified intervals for the sequence and in this example, it is used for illustrating the positions of the protein domains with cross-link features that enable users to locate the website or page of the original database or article. **(B)** The notification part shows any type of defined modifications as a red asterisk above the sequence per position and displays the searched position in a species above the alignments. **(C)** The sequence alignment part contains the imported alignment data with the previously selected color scheme. Also, rounded (circle) positions indicate that at least one genetic variation or modification exists in this position. A rectangular white background pop-up box appears when the mouse hovers the specific position in the sequence and the genetic variations and modifications are listed in this pop-up box. On the bottom, an auto-generated ‘Consensus’ sequence is displayed. On the left side, species names contain cross-reference links for referring to the dedicated page of the sequence according to its protein identifier such as a UniProt number and the near-white ‘x’ button enables users to hide the sequence from the alignment together with its identifier. **(D)** A position in the sequence of any species listed in the alignment can be searched and the sequence alignment data in FASTA format can be downloaded with the blue button and visualization of alignment data can be exported as PNG format. Also, with the green ‘Reset’ button, it is available to reload the viewer.

MSABrowser introduces four major novelties: first, the pliable annotation of genetic variants (c.88C>G or p.Pro30Ala), orthologous variants (OrthoVars), or posttranslational modifications (PTMs) (ubiquitination at Lysine 2563; K2563-ub) into the respective sequence positions on the PSA and MSA **(Fig. 2A and B**) (Pir *et al*., 2021); second, multiple annotations, such as protein domains (SH3 domains) and/or user-specified intervals can be added at the same time to the corresponding positions; third, the variant-specific annotations, including phenotypic data, variant ID, and allele frequency, can be integrated into the corresponding positions. For example, p.R79Q in ARL13B (Protein ID = NP_001167621.1) has several variant-specific annotations, including variant ID (rs121912606), an allele frequency (3.98e-6), predicted as a pathogenic variant, and disease association (causing Joubert syndrome) (Karczewski *et al*., 2020; Cantagrel *et al*., 2008), and all of these annotations can easily be co-viewed at the respective sites. Finally, scrolling through PSAs/MSAs, searching, and custom styling are implemented. While the MSABrowser can easily integrate annotations (OrthoVars, PTMs, allele frequency, variants, and variant ID, etc.) into the corresponding positions, other MSA tools lack position specific annotation integration features **(Fig. 3**).

**Fig. 2.**
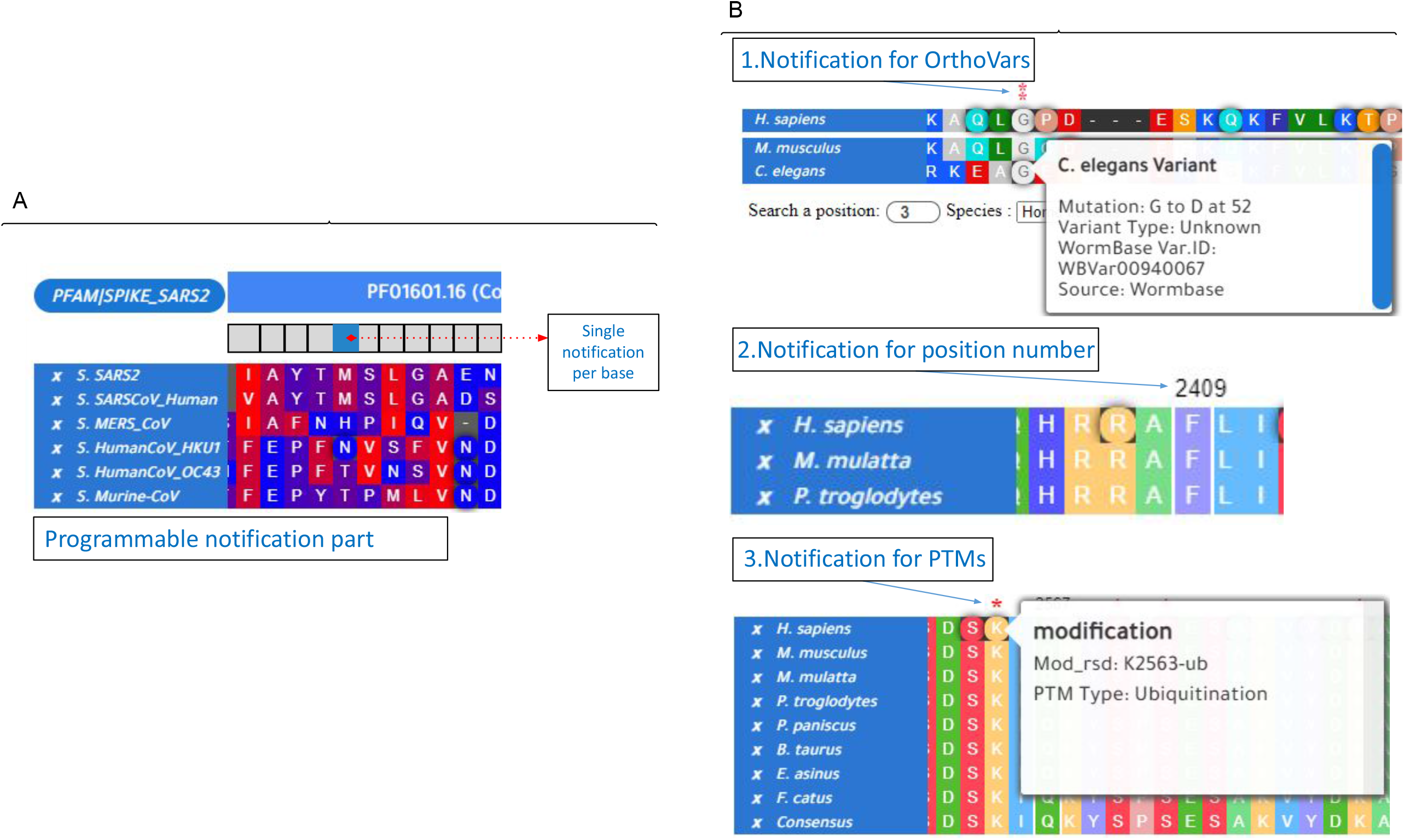
Example demonstration with the MSABrowser tool. **(A)** Visualization of multiple sequence alignments of six virus spike proteins with the MSABrowser tool. The positions with the annotations are marked in a circle, while the positions without annotations are displayed in a square. The full MSA comparisons with annotations can be found at our dedicated GitHub site http://thekaplanlab.github.io/ (**B)** Shown is the display of orthologous variants (OrthoVars), the positions of amino acid position or nucleotide, or posttranslational modifications (PTMs) with the programmable notification part of MSABrowser.

**Fig. 3.**
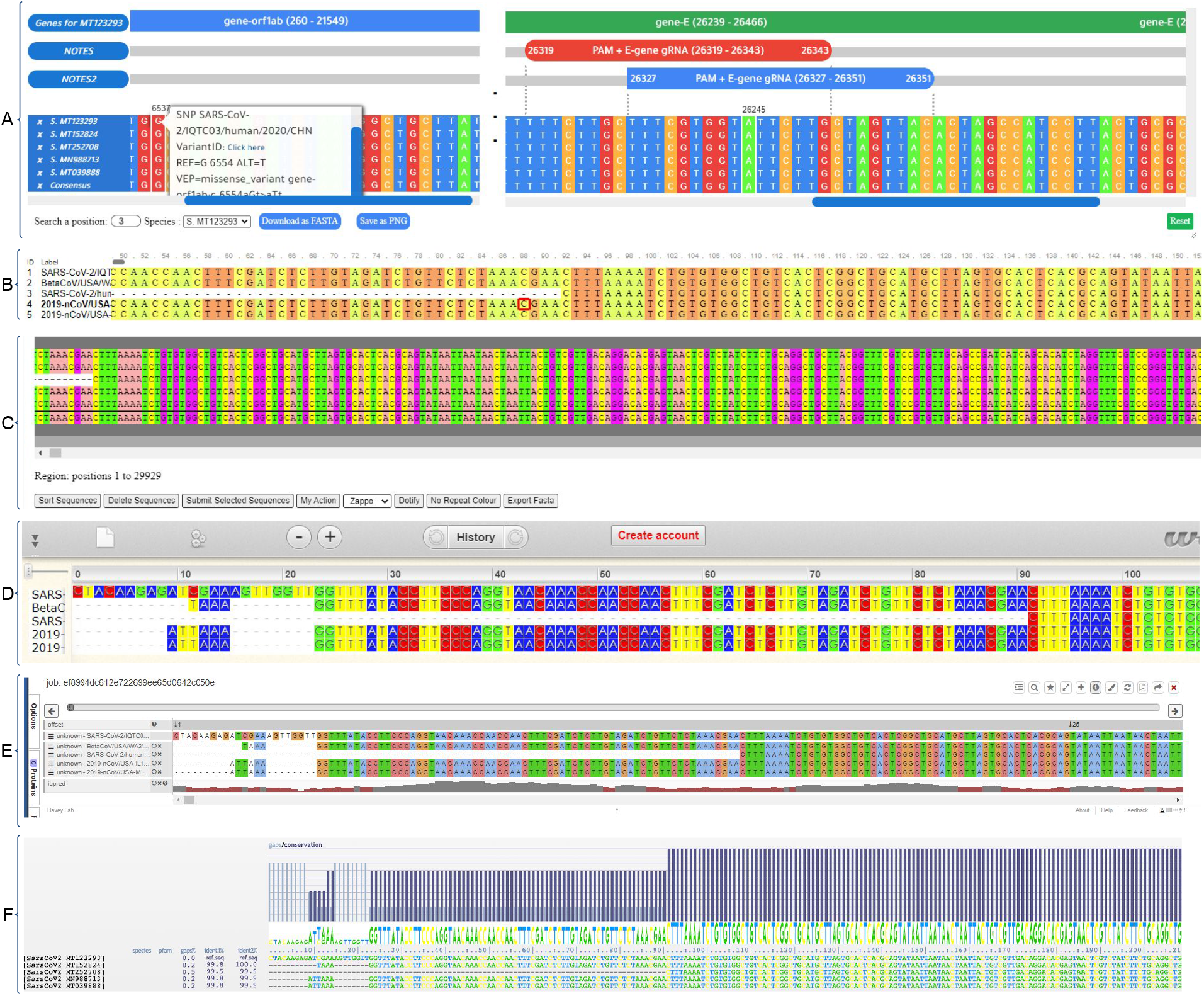
Comparison of MSA visualizers. The multiple sequence alignment of different genomes of the severe acute respiratory syndrome coronavirus 2 isolates (SARS-CoV-2: MT123293, MT152824, MT252708, MN988713, and MT039888) was created, followed by visualization with separate MSA viewers. **(A)** Shown is the MSA visualization with MSABrowser, which enables addition of annotations (e.g. domains and notes) on top of the MSA. MSABrowser allows users to incorporate variant-specific annotations (missense variation, disease associations, variant ID, allele frequency, etc.). A popup will show up when users can click on the circled amino acid or nucleotide position to display the annotations. Shown is a missense variation (G6537T) in SARS-CoV, an example of a particular annotation of a nucleotide position, MSABrowser enables users to remove the desired sequence by clicking the X button which appears in the far left of each line. Users can look up positions, download the FASTA and save the MSA as PNG. **(B)** Shown is MSAViewer tool on the same alignment as in **A**. Users can scroll to left and right to see the rest of MSA. When the user clicks on a position, the amino acid is highlighted with a red square as in the position 88. **(C)** Shown is JSAV. It is possible to sort and delete sequences, add new sequences, change the color schema, and export FASTA with the buttons listed below the MSA. **(D)** Shown is Wasabi in which zoom in and zoom out options are enabled and scrolling is necessary to see the rest of the sequence. **(E)** Shows Proviz where users are able to search for a motif, switch to full screen, export the MSA and share it as a URL using the buttons located in the top right corner. **(F)** Shown is AlignmentViewer. For each sequence in the alignment, gaps ratio, and identification ratio to the reference sequence is provided. Gaps and conservation per position is also shown above the MSA.

## 2 Availability and implementation

PSAs and MSAs are the fundamental methods for the alignment of any sequences of DNA, RNA, and protein (Chenna, 2003; Higgins and Sharp, 1988). The MSABrowser imports PSA and MSA data in FASTA format with a file, and variations and sequence annotation data in JavaScript Object Notation (JSON) (Pearson, 1999). After parsing the alignment data and creating the consensus sequence, it then creates two main components: the annotation part and the sequence alignment part. For performance purposes, instead of rendering all the alignment data at once, the MSABrowser renders as the user navigates through the sequence alignment. The positions consisting of the modifications such as post-translational modifications or variations are highlighted with shadow or asterisk together with rounded boxes on the corresponding positions of nucleotide or amino acids and hovering on them triggers a pop-up that shows the details of variations and modifications or any other provided notes for the position.

The MSABrowser has multiple ways of navigating the alignment. Firstly, by scrolling through the sequence alignment and secondly, by specifying either amino acid or nucleotide position and the species in the bottom panel. Users can hide sequences from the alignment by selection. Additionally, a cross-reference link is automatically generated based on the sequence identifiers from the imported FASTA file. Therefore, users may click the species names to jump to the sequence database (i.e. Ensembl, NCBI, and UniProt). For visualizing the alignments, users might choose between 13 predefined color schemes. The MSABrowser is capable of exporting alignment as a FASTA file format and the visualization as a publication-quality figure in Portable Network Graphics (PNG). Furthermore, the detailed comparison of features among other web-based visualization tools (Veidenberg *et al*., 2016; Yachdav *et al*., 2016; Martin, 2014; Larsson, 2014; Jehl *et al*., 2016; Hossain, 2019) is available in Supplementary Table 1.

## 3 Conclusion

MSABrowser is the most recently created tool that allows the visualization of MSAs, genetic variations, posttranslational modifications, and protein domains at the same time. MSABrowser makes it much easier to display orthologous variants between different species (Pir *et al*., 2021). Importantly, it does not require the installation of any software as it runs on any modern browser that is pre-installed on computers. Its portability, speed, and recently updated date make it a viable unified sequence alignment, variations, and annotations visualization tool.

## 4 Acknowledgements

We thank Sebiha Cevik for comments on the manuscript.

## Code availability

Codes and dataset used for creating Figure 1 are available from https://github.com/thekaplanlab/msabrowser

